# SUGP1 loss is the sole driver of SF3B1 hotspot mutant missplicing in cancer

**DOI:** 10.1101/2025.02.17.638713

**Authors:** Peiqi Xing, Pedro Bak-Gordon, Jindou Xie, Jian Zhang, Zhaoqi Liu, James L. Manley

**Author notes:** P.X., P.B-G., J.X., and J.Z. contributed equally to this work.

## Abstract

SF3B1 is the most frequently mutated splicing factor in cancer. Mechanistically, such mutations cause missplicing by promoting aberrant 3’ splice site usage; however, how this occurs remains controversial. To address this issue, we employed a computational screen of 600 splicing-related proteins to identify those whose reduced expression recapitulated mutant SF3B1 splicing dysregulation. Strikingly, our analysis revealed only two proteins whose loss reproduced this effect. Extending our previous findings, loss of the G-patch protein SUGP1 recapitulated almost all splicing defects induced by *SF3B1* hotspot mutations. Unexpectedly, loss of the RNA helicase Aquarius (AQR) reproduced ∼40% of these defects. However, we found that AQR knockdown caused significant *SUGP1* missplicing and reduced protein levels, suggesting that AQR loss reproduced mutant SF3B1 splicing defects only indirectly. This study advances our understanding of missplicing caused by oncogenic *SF3B1* mutations, and highlights the fundamental role of SUGP1 in this process.

## Introduction

RNA splicing entails a two-step reaction by which non-coding introns are excised from precursor messenger RNAs (pre-mRNAs) and the remaining exons are ligated to generate mature mRNA transcripts^1^. More than 95% of human transcripts are subject to the process of alternative splicing, whereby pre-mRNAs are differentially spliced to generate multiple proteins with functional diversity^2,3^. Splicing occurs in a large and dynamic structure called the spliceosome, consisting of over 100 proteins and five small nuclear RNAs (snRNAs) assembled into snRNPs^4^. As a fundamental cellular process, dysregulation of mRNA splicing has been implicated in multiple diseases including cancer, in which it can contribute to tumor initiation, progression and treatment resistance^5^.

Multiple genes encoding splicing factors are recurrently mutated in cancer, highlighting splicing dysregulation as a genetic driver of oncogenesis^6^. Amongst these, mutations in the gene encoding the core splicing factor SF3B1 are the most common, frequently occurring as “hotspot” heterozygous point mutations affecting specific amino acid residues concentrated in the so-called HEAT repeat domain (SF3B1^HEAT^), specifically repeats H4–H7^7–9^. *SF3B1* hotspot mutations promote the recognition of aberrant branchpoints, resulting in increased usage of cryptic 3’ splice sites (3’ss) typically located 10-30nt upstream from canonical 3’ss^10–12^. While other types of missplicing have been reported^13^, these constitute a minority, and cryptic 3’ss are the most prevalent. This change in splicing is consistent with the role of SF3B1 as a core component of the U2 small nuclear ribonucleoprotein (snRNP) complex, which is responsible for branch site (BS) recognition and spliceosome assembly during the early stages of splicing^10^. Functional studies of the downstream splicing defects arising in *SF3B1*-mutant cancers, including myelodysplastic syndromes (MDS)^14–16^, chronic lymphocytic leukemia (CLL)^17^, uveal melanoma (UVM)^13^, breast^18^ and pancreatic cancer^19^, have elucidated specific aberrant splicing events critical for the establishment and maintenance of such cancers. However, the precise mechanism(s) by which *SF3B1* mutations lead to cryptic alternative 3’ss selection and the key interacting partner proteins involved in this process remains controversial.

Previous systematic analyses of RNA splicing profiles across The Cancer Genome Atlas (TCGA) database revealed that somatic mutations in *SUGP1* (SURP and G-patch domain containing 1) mimic the mutant *SF3B1* splicing dysregulation^20,21^. Importantly, these findings are consistent with earlier studies showing that *SF3B1* hotspot mutations disrupt an interaction between SF3B1 and SUGP1 that is necessary for correct BS recognition during splicing, and that SUGP1 knockdown can recapitulate splicing defects observed in *SF3B1* mutant cells^22^. Subsequently, the regions of *SUGP1* encompassing the previously identified cancer mutations, which flank its G-patch domain, were found to interact directly with SF3B1 HEAT repeats harboring cancer-associated hotspot mutations, resulting in a conformational change of SUGP1 that exposes its G-patch domain to interact with and thereby activate the DEAH box helicase DHX15^23,24^. Thus, cancer-associated hotspot mutations in SF3B1 or SUGP1 prevent activation of DHX15, which in turn leads to aberrant use of upstream branch points and cryptic 3’ss characteristic of tumors with mutant SF3B1. However, whether this mechanism entirely explains mutant SF3B1 missplicing remains unclear.

Recent studies have suggested that a number of splicing factors, lacking known mutations in cancer, are potentially linked to mutant *SF3B1* splicing dysregulation. For example, Benbarche et al. found that the G-patch domain-containing protein GPATCH8 can exert an antagonistic effect on SUGP1 by competing with SUGP1 for the same binding region of DHX15. Silencing GPATCH8 enhanced the DHX15/SUGP1 interaction and corrected one-third of mutant SF3B1 missplicing defects^25^. In addition, two members of the DEAD-box ATPase family, DDX42 and DDX46 (also known as PRP5), have been reported to compete for binding to SF3B1 at the hotspot region^26,27^, and *SF3B1* cancer mutations were found to weaken these interactions^28^. However, knockdown (KD) of DDX46 did not induce cryptic 3′ss usage in any of the top targets of mutant SF3B1^24^. Notwithstanding, it remains to be determined if loss of DDX42/46, or in fact DHX15, can also recapitulate the missplicing induced by *SF3B1* mutations, as SUGP1 loss or mutation does.

As indicated above, computational screening of splicing changes has contributed to a better understanding of mutant SF3B1 splicing dysregulation. Even though TCGA offers rich genomic information on numerous tumor types, instances of mutations that can mimic mutant SF3B1 splicing defects are in general rare^21^. Beyond mutations, altered expression levels of RNA-binding proteins (RBPs) can also contribute to cancer-associated splicing abnormalities^29–31^. Thus, further computational screening may reveal additional RBPs/splicing factors whose expression changes reproduce the aberrant splicing patterns induced by *SF3B1* mutations, thereby providing new clues for fully elucidating the mechanism underlying mutant SF3B1 missplicing.

In this study, we first employed a computational approach to define a high-confidence list of cryptic 3’ss events characteristic of *SF3B1* mutations across different cancer types and cell lines. Based on these events, we conducted a comprehensive screen to evaluate whether RBP/splicing factor loss reproduced the splicing defects induced by *SF3B1* mutations. Our findings identify loss of SUGP1 as the sole driver of SF3B1 mutant misplacing, underscoring the indispensable role of SUGP1 in SF3B1 function during BS recognition and 3’ss selection that is perturbed by *SF3B1* hotspot mutations in cancer.

## Results

### Identification of a comprehensive high-confidence list of cryptic 3’ splice sites used in *SF3B1*-mutant cells

As discussed above, multiple studies have shown that *SF3B1* hotspot mutations typically result in missplicing arising from the use of cryptic 3’ss in affected transcripts. These analyses used RNA from a variety of sources, ranging from patient samples to CRISPR-generated cell lines, and identified hundreds to thousands of transcripts, depending on technical parameters^5^. Although several common targets have been identified and validated, the differences in biological samples and computational analyses have prevented the establishment of a comprehensive list of high-confidence mutant SF3B1 targets. Such a list would be of value for many studies, including our efforts here to identify splicing factors whose function might be relevant to mutant SF3B1-induced missplicing.

To achieve the above goals, we designed a computational framework to identify and quantify cryptic 3’ss utilization using 63 samples containing *SF3B1* mutations (SF3B1^MUT^) and 56 *SF3B1* wild-type controls (SF3B1^WT^) from six different datasets, including for example MDS, CLL and UVM (Figure 1*A*, Table S1 and Methods). We first identified a total pool of 22,158 cryptic 3’ss candidates with at least 200 cryptic read counts across all SF3B1^MUT^ samples. Next, to capture authentic 3’ss changes resulting from *SF3B1* mutations, we compared the Percent-Spliced-In (PSI) values of each event between SF3B1^MUT^ and SF3B1^WT^. Cryptic events associated with a *p*-value <0.05 (t-test) and a ΔPSI >0.1 in three or more of the six cohorts and with a distance of 3-100 nt between cryptic and canonical 3′ss were selected (n=316). Lastly, we manually examined every event using Integrative Genomics Viewer (IGV). This analysis identified 295 cryptic 3’ss events as high-confidence recurrent splicing changes arising from *SF3B1* mutations (Table S2). Additionally, we also generated a “core” list of 123 cryptic 3’ss events that met the above criteria across all six cohorts (Table S3). We also validated the specificity of the 295 cryptic 3’ss events by unsupervised clustering and principal component analysis (PCA), both of which clearly distinguished SF3B1^MUT^ from SF3B1^WT^ (Figure S1*A, B*). As expected, and consistent with previous studies^10–12^, a large majority of the cryptic 3’ss were located 10 to 30 nt upstream of the respective canonical 3’ss (Figure S1*C*), although a small fraction were located further upstream and, notably, 21% were located downstream.

**Fig 1.**
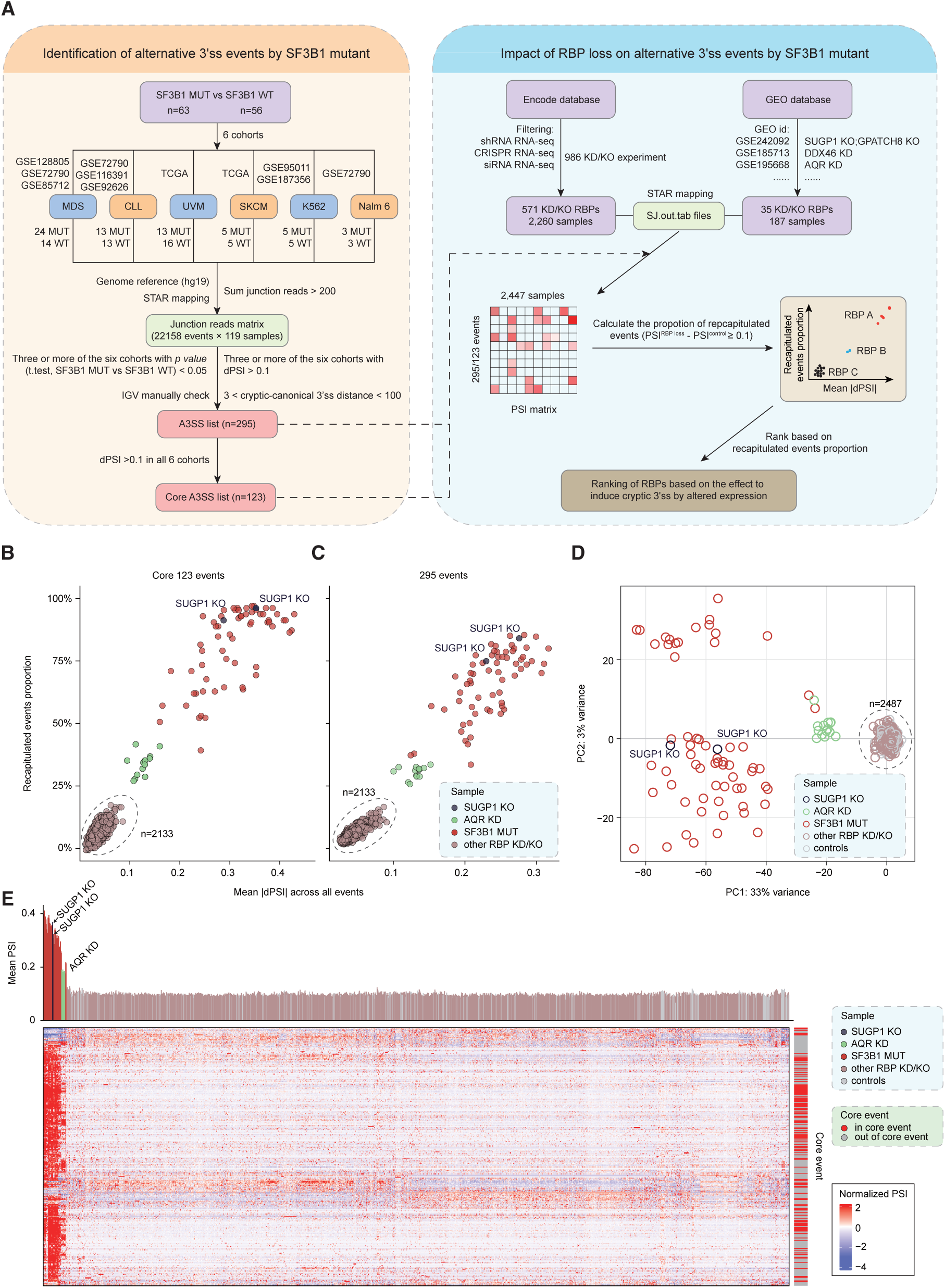
Identification of SF/RBPs whose expression loss recapitulates cryptic 3’ss usage by *SF3B1* hotspot mutations. (*A*) Schema of the computational pipeline used to identify cryptic 3’ss evenst induced by *SF3B1* mutations (left) and to evaluate effects of SF/RBP loss on cryptic 3’ss events induced by *SF3B1* mutations (right) (see details in Methods). (B and C) Scatter plot representations of SF/RBPs whose expression loss positively correlated with cryptic 3′ss usage. The horizontal axis shows the averaged absolute PSI change values between SF/RBP KD/KO samples and controls, and the vertical axis shows the percentage of recapitulated events. Analysis was performed on the 123 core (*B*) and 295 (*C*) SF3B1^MUT^-specific events. (*D*) Principal component analysis (PCA) projection of all RNASeq samples using the PSI values of 295 cryptic 3’ss events. (*E*) Hierarchical clustering using Euclidean distance and heatmap analysis of the usage of 295 SF3B1^MUT^-specific events in all SF/RBP KD/KO, SF3B1^MUT^ and control samples. Rows represent the 295 SF3B1^MUT^-specific events, while columns represent the samples. Matrix values correspond to raw PSI values. Each vertical bar represents the mean of the PSI values of the 295 events in each of the SF/RBP loss and control samples. The row annotation bar plot on the right of the heatmap indicates whether the event is in the core 123 event list.

To characterize the cryptic 3’ss further, we performed sequence motif analysis on the nucleotide sequences ±50bp from both cryptic and associated canonical 3’ss of the 295 events, using 500 randomly selected canonical 3’ss without cryptic events as controls. (Figure S1*D*, Methods). Consistent with previous observations^11,14,17^, a relatively short and weak polypyrimidine tract interrupted with adenosines was found upstream of cryptic 3’ss compared to canonical and control sites (Figure S1*D*). Altogether, we have identified robust lists of cryptic 3’ss that are highly specific to and characteristic of SF3B1^MUT^ cells.

### Depletion of only two splicing-related proteins recapitulates SF3B1^MUT^ missplicing

We next used the above lists of cryptic 3’ss induced by mutant SF3B1 as benchmarks to assess the ability of splicing factor (SF)/RBP loss to reproduce the splicing dysregulation brought about by *SF3B1* mutations. To this end, we collected RNA-seq data on 600 SF/RBP KD/KO samples from ENCODE and GEO databases (Table S4). We then calculated the percentage of cryptic 3’ss that could be recapitulated by SF/RBP loss (Figure 1*A*, Methods, Table S5) and ranked all SF/RBPs based on the percentage of recapitulated events (Table S6). As a positive control, we also performed this analysis on each of the SF3B1^MUT^ samples used to generate the event list. We found that all of these SF3B1^MUT^ samples consistently captured most cryptic 3’ss events, with percentages varying from 40% to 98% for the core list and 34% to 85% for the 295 event list (Figure 1*B, C*). The variation could be due to multiple factors, including the different SF3B1-mutated residues, which are known to produce distinctive splicing patterns^32^, the variant allele frequency, the affected cancer types as well as variations between individual samples. This observation further supports our strategy of utilizing as many as possible SF3B1^MUT^ samples for generation of the lists of cryptic 3’ss events that occur across different cohorts. Notably, SF3B1 KO/KD did not significantly recapitulate these splicing defects (Table S6), consistent with the fact that *SF3B1* mutations are neomorphic change-of-function mutations rather than loss-of-function^32^.

Among the SF/RBP samples, exceeding 2,000 in total, KD/KO of only two spliceosomal proteins recapitulated the SF3B1^MUT^ events to a significant extent. Strikingly, both SUGP1 KO samples reproduced a high percentage of the splicing defects, ∼95% of the 123 core events and ∼80% of the 295 events. Notably, these values were very similar to the top-ranked control SF3B1^MUT^ samples (Figure 1*B, C*), indicating that SUGP1 loss underlies all or nearly all SF3B1^MUT^ missplicing. The second SF/RBP that recapitulated SF3B1^MUT^-induced missplicing significantly when depleted was unexpected, as KD of the RNA helicase Aquarius (AQR) was found to reproduce 28% to 42% for the 123 event list (Figure 1B), and 26% to 34% for the 295 event list (Figure 1C and Table S6). We also applied unsupervised PCA to the 295 event list across all SF/RBP loss, SF3B1^MUT^ and SF3B1^WT^ samples, which gave a similar conclusion (Figure 1*D*). In addition, to obtain a global view of missplicing of the 295 3’ss events, we performed unsupervised hierarchical clustering on all the samples (Fig. 1*E*). Overall, *SF3B1* mutations, SUGP1 KO and AQR KD samples were clustered closer together and with significantly higher PSI than observed with all other SF/RBP KD/KO and WT samples (Fig. 1*E*), indicative of the similarity between these samples in cryptic 3’ss usage.

### AQR loss recapitulates SF3B1^MUT^-specific cryptic 3’ss events by downregulating SUGP1

As mentioned above, our previous studies found that cancer-associated mutations in *SUGP1,* as well as SUGP1 KD, mimic the missplicing caused by *SF3B1* mutations^20,21^. Although others have suggested alternate mechanisms (see above), it was thus not entirely unexpected that our analysis revealed the striking overlap between SUGP1 loss and *SF3B1* mutations in inducing cryptic 3’ss use. However, it was completely unanticipated that AQR, a protein with limited evidence of involvement in BS recognition^33^ (see below), was the only other SF/RBP whose loss mimicked SF3B1^MUT^ missplicing (Figure 1*B-E*). To validate this finding, we first used unsupervised hierarchical clustering and PCA with AQR KD samples from K562 and HEK293T cells, which revealed that AQR loss recapitulated 39% of SF3B1^MUT^-specific cryptic 3’ss in the 123 event core list and 32% in the 295 event list (Figure 2*A, B* and Figure S2*A-C*). We next examined experimentally whether AQR KD induced SF3B1^MUT^ cryptic 3’ss use, with a panel of top target transcripts identified above as examples (Figure 2C). To do this, we used two independent siRNAs to deplete AQR in K562 cells (KD efficiencies shown in Figure 2D). RT-PCR showed that AQR KD indeed robustly induced use of cryptic 3’ss, recapitulating the splicing pattern we observed in SF3B1^K700E^ K562 cells^22^ (Figure 2*E*).

**Fig. 2.**
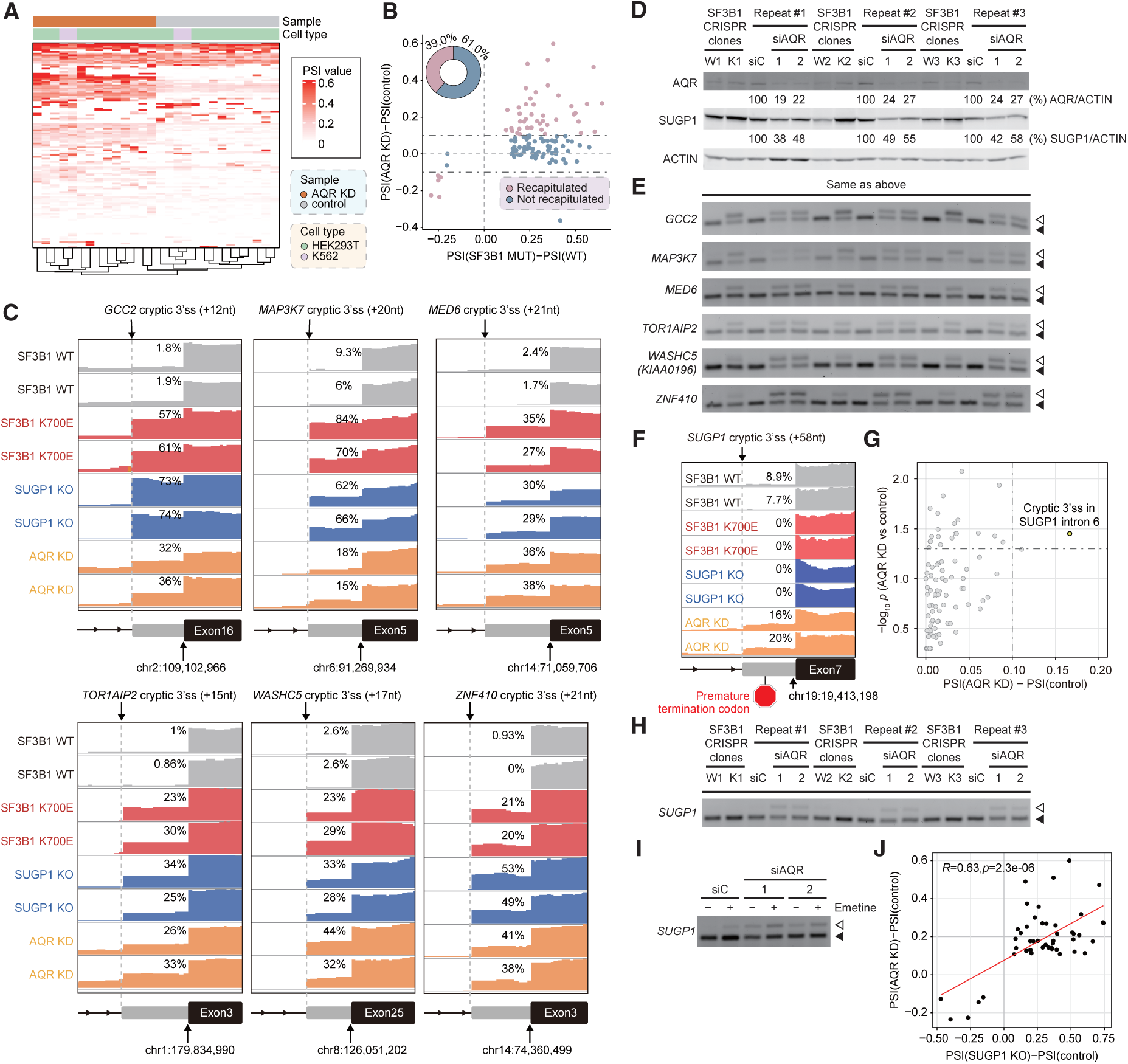
AQR KD recapitulates ∼40% of SF3B1^MUT^-specific cryptic 3’ss events and induces *SUGP1* mRNA missplicing. (*A*) Hierarchical clustering and heatmap analysis of the usage of 123 core events in AQR KD samples and corresponding control samples. Rows represent the 123 core events, while columns represent the samples. (*B*) Scatter plot displaying the ΔPSI values for the 123 core events in AQR KD samples compared with SF3B1^MUT^ samples. The horizontal axis shows the ΔPSI value between SF3B1^MUT^ and control samples, while the vertical axis shows the ΔPSI value between AQR KD sample and control sample. The pie plot (upper-left) indicates the proportion of 3’ss events recapitulated (brown) or not recapitulated (blue) by AQR KD. (*C*) IGV plots of cryptic 3’ss event recapitulated by AQR loss in samples with SF3B1 mutations, SUGP1 loss, AQR loss or WT. PSI values are labeled for each track. (*D*) Western blotting of protein extracts for SUGP1 from either CRISPR-engineered K562 cells with WT (W) or K700E (K) SF3B1, or parental K562 cells electroporated with a negative control siRNA (siC) or one of two independent siRNAs targeting AQR. (*E*) Total RNA was extracted from the cells as in (*D*), followed by RT-PCR to identify the cryptic 3′ss and associated canonical 3′ss produced from splicing of the transcripts indicated. (*F*) IGV plots of cryptic 3’ss usage in *SUGP1* intron 6 in WT, SF3B1^K700E^, SUGP1 KO and AQR KD samples. PSI values are shown for each track. (*G*) Scatter plot representation of differentially spliced 3’ss changes on U2 complex genes between AQR KD and controls showing the magnitude (difference of PSI; x-axis) and significance (-log10(q-value); y-axis). (*H*) Same RT-PCR procedure as in (*E*) except to detect cryptic 3’ss usage in *SUGP1* intron 6. (*I*) K562 cells were electroporated with a negative control siRNA (siC) or one of two AQR siRNAs as indicated, treated with (+) or without (−) emetine, followed by PT-PCR of isolated RNA to detect the cryptic 3′ss and associated canonical 3′ss produced from splicing of *SUGP1* intron 6. (*J*) Scatter plot depicting the correlation between AQR KD and SUGP1 KO on the SF3B1^MUT^-specific events recapitulated by AQR KD. The horizontal axis shows the PSI changes between SUGP1 KO and control samples, while the vertical axis shows the PSI changes between AQR KD and control samples. The indicated R value and *p-*value were determined by Pearson correlation.

Previous studies have provided evidence that AQR KD can lead to over 30,000 altered splicing events^34,35^. Indeed, we observed that over 2,000 cryptic 3’ss were utilized following AQR KD (Table S7; see also Figure S3A below). This raised the possibility that the ∼40% of SF3B1^MUT^-missplicing events recapitulated by AQR KD arose only by coincidence, reflecting chance overlap. If this were in fact the case, then a similar degree of overlap would be expected to occur with any other samples in which splicing has been disrupted. To test this, we utilized RNA-seq data from CRISPR-engineered K562 cells with hotspot mutations in U2AF1 (three cases of S34F vs. three WTs) and SRSF2 (four cases of P95H vs. four WTs), which are the two most commonly mutated splicing factor genes in cancer after *SF3B1*^36^ (Table S8). We first identified the top 295 events that were most specific to the SRSF2 or U2AF1 mutant samples (Figure S2*D, E* and Methods), and then compared the correlation between the PSI changes of the 295 events from all three spliceosomal mutations and *AQR* KD. The results showed that the splicing changes of AQR KD significantly correlated only with the splicing defects detected in the SF3B1^MUT^ samples (Figure S2*F*). These findings indicate that the overlap we detected between AQR KD and SF3B1^MUT^ samples was specific.

Since AQR has not been shown to function directly with SF3B1 during BS recognition, we wondered whether AQR loss recapitulated SF3B1^MUT^ splicing defects indirectly. An interesting possibility was that this might have resulted from downregulation of SUGP1. To test this hypothesis, we performed western blot (WB) analysis of SUGP1 with AQR KD cells and indeed found a significant decrease (∼50-60%) of SUGP1 protein levels upon AQR KD (*p*-value <0.0001 for both siAQR) (Figure 2*D* and Figure S2*G*). One explanation for this, consistent with the massive missplicing induced by AQR KD, is that *SUGP1* transcripts were misspliced, resulting in reduced levels of *SUGP1* mRNA. Using the same computational pipeline for 3’ss event identification as above, we examined the splicing changes in transcripts encoding U2 complex-associated splicing factors annotated in the UniProt database^37^. Strikingly, the top target we identified in AQR KD cells was a cryptic 3’ss in *SUGP1* intron 6, usage of which led to insertion of a 58nt intronic sequence in the mRNA that introduces a premature termination codon (Figure 2*F-*H). The misspliced *SUGP1* mRNA was subject to nonsense mediated decay (NMD), as the RT-PCR products encompassing the cryptic 3’ss were increased upon NMD inhibition by emetine^37^ (Figure 2*I*). Notably, AQR KD induced use of this novel cryptic 3’ss by a mechanism different from *SF3B1* mutations, as *SF3B1* mutations did not induce this event (Figure 2F). Finally, consistent with the above, we found a significant positive correlation of ΔPSI between SUGP1 loss and AQR loss on AQR KD-recapitulated SF3B1^MUT^ 3’ss events, which accounted for 32.2% of the 295 events (Figure 2*J*).

In addition to *SUGP1*, multiple transcripts misspliced following AQR KD are involved in RNA processing. One example is *TFIP11* (Figure S3*A*), which encodes a G-patch protein that like SUGP1 functions in splicing by activating DHX15^38^. Interestingly, another is the transcript encoding Senataxin (SETX), an RNA:DNA helicase that functions to resolve cotranscriptional R loops^39^. *SETX* was in fact one of the most misspliced transcripts following AQR KD, with >40% of transcripts using a cryptic 3’ss in intron 23 (Figure S*3B*). This missplicing, which was not observed in SF3B1^MUT^ or SUGP1 KD cells, generated a PTC, and as shown by WB reduced protein levels strikingly, by ∼80% (*p*-value = 0.0002 for siAQR-1 and 0.0001 for siAQR-2) (Figure S3*C, D*). Notably, it has been proposed that AQR is directly involved in resolving R loops, as the abundance of these structures is increased upon AQR KD^40–43^. However, AQR has only been shown to possess helicase activity capable of unwinding RNA:RNA hybrids^33^, and is thought to be localized exclusively in spliceosomal complexes^35^. Our data thus suggest that AQR loss leads only indirectly to an increase in R loops, due to SETX downregulation.

### SUGP1 is the only G-patch protein whose loss recapitulates splicing defects induced by SF3B1^MUT^

A recent report found that another G patch-containing protein, GPATCH8, is required for a subset of the missplicing events induced by *SF3B1* mutations. GPATCH8 was found to exert an antagonistic effect on SUGP1, and silencing of GPATCH8 in *SF3B1*-mutated cells corrected one-third of SF3B1^MUT^-dependent splicing defects^25^. Thus, we next sought to compare the effect of SUGP1 or GPATCH8 loss on the cryptic 3’ss changes observed in SF3B1^MUT^ cells. We compared the PSI distributions of these events along an axis of SF3B1^WT^, SF3B1^MUT^ and SUGP1 KO in SF3B1^MUT^ cells. Interestingly, we observed a monotonic increase of the overall PSI values along this axis, in which SUGP1 loss in SF3B1^MUT^ cells led to further increased 3′ss usage in about 85% of the 123 core events (Figure 3*A, B*) and 78% of the 295 events (Figure S4*A, B*) compared to *SF3B1* mutation alone (Table S9). Consistent with results reported in the previous study^28^, this phenomenon was opposite to that observed with GPATCH8 KO. Compared to *SF3B1* mutation alone, GPATCH8 KO in SF3B1^MUT^ cells led to decreased 3′ss usage in about 64% of the 123 core events (Figure 3*C, D*) and 66% of the 295 events (Figure S4*C, D*). This rescue proportion was higher than the ∼30% reported by Benbarche et al. upon GPATCH8 KO^25^. A likely reason for this discrepancy is that we selected the most prominent splicing events from different SF3B1^MUT^ datasets, including only cryptic 3’ss events, which are not only the most common type of splicing change in SF3B1^MUT^ cells but also the most likely to reflect direct effects.

**Fig. 3.**
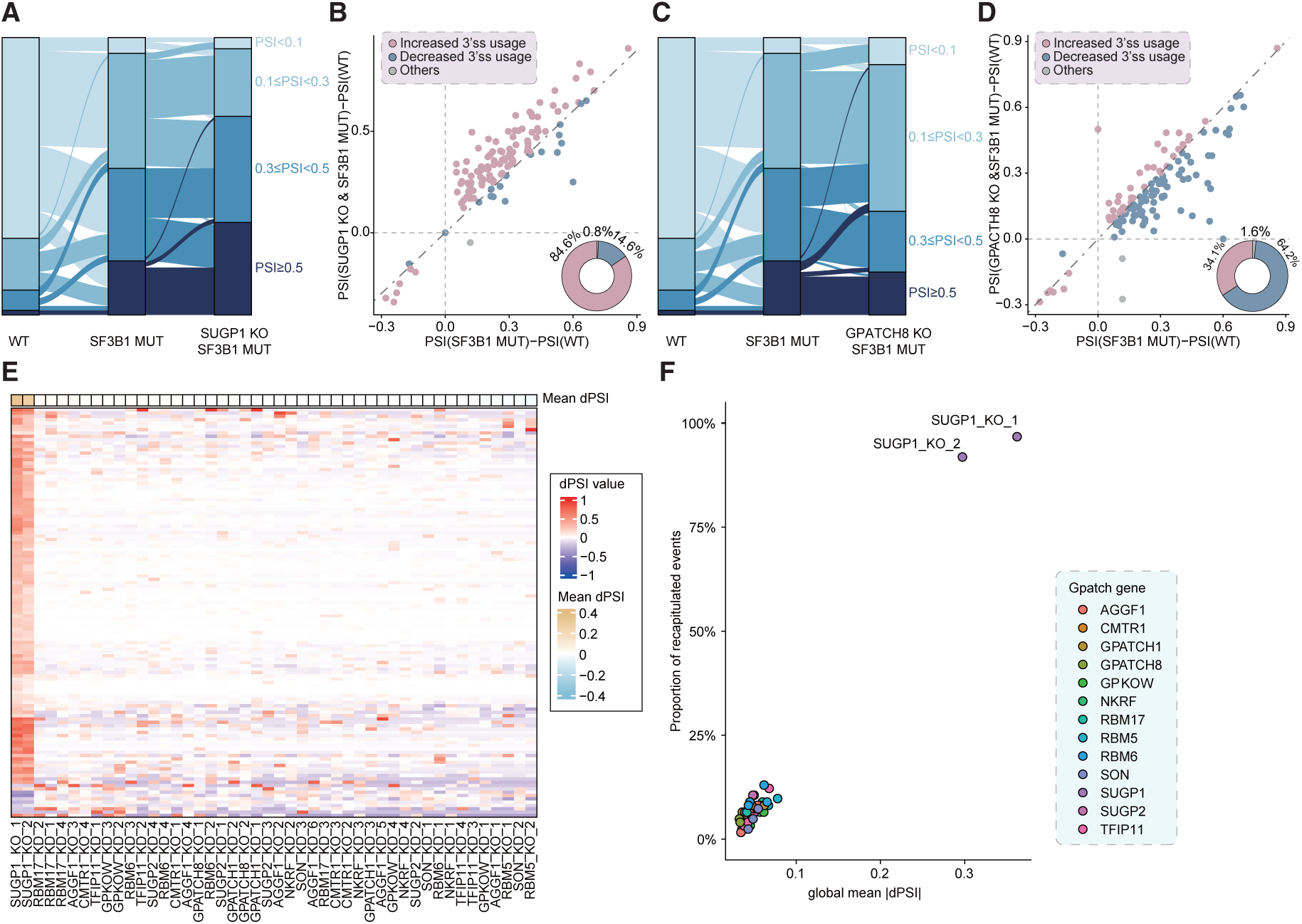
Distinct effects of G-patch protein loss on SF3B1^MUT^-specific cryptic 3’ss events. (*A*) Sankey diagram illustrating the changes in PSI values of 123 core events induced by *SF3B1* mutation along an axis of SF3B1^WT^, SF3B1^MUT^, and SUGP1 KO in SF3B1^MUT^ cells. Different colors represent the range of PSI values. (*B*) Scatter plot depicts ΔPSI values of 123 core events in SF3B1^MUT^ cells compared with SF3B1^MUT^ and SUGP1 KO cells. The horizontal axis shows the ΔPSI value between SF3B1^MUT^ and control samples, while the vertical axis shows the ΔPSI value between SF3B1^MUT^ and SUGP1 KO and control samples. Pie plot (lower-right) indicates proportions of increasingly (brown) and decreasingly (blue) used cryptic 3’ss in SF3B1^MUT^ and SUGP1 KO samples compared to SF3B1^MUT^ alone sample. (*C*) Similar to (*A*) but for GPATCH8. (*D*) Similar to (*B*) but for GPATCH8. (*E*) Hierarchical clustering and heatmap analysis of the ΔPSI values of 123 core events in samples with KO/KD of G-patch protein-encoding genes. Rows represent the 123 core events, while columns represent the samples. The column annotation bar plot at the top of the heatmap indicates the averaged ΔPSI values across the 123 core events. (*F*) Scatter plot representations of G patch-containing proteins (Gpatch gene) whose expression loss positively correlated with cryptic 3′ss usage. The horizontal axis shows the average ΔPSI values between G-patch protein KD/KO samples and controls, and the vertical axis shows the percentage of recapitulated events. Analysis was performed on the 123 core events.

Given that DHX helicases such as DHX15 can be activated by multiple G patch-containing proteins^44^, we wondered if loss of any G-PATCH protein other than SUGP1 might also mimic SF3B1^MUT^ missplicing. To examine this possibility, we analyzed splicing patterns induced by loss of all G patch-containing RBPs for which appropriate RNA-seq data was available, and found that SUGP1 was the only one whose loss recapitulated the splicing changes induced by mutant SF3B1^MUT^ (Figure 3*E, F* for the 123 core events, Figure S4*E, F* for 295 events).

### Loss of DHX15 but not DDX46 or DDX42 recapitulates SF3B1^MUT^ missplicing

As mentioned above, two DDX-type RNA helicases, DDX46 and DDX42, are known to associate with human U2 snRNP, and to interact with SF3B1 at the region where hotspot mutations occur^26–28^. Given that *SF3B1* cancer mutations can disrupt the DDX46 interaction, it has been suggested that DDX46 loss from the spliceosome may underlie SF3B1^MUT^-induced missplicing^31^. However, we previously demonstrated experimentally that DDX46 KD did not recapitulate missplicing of any of a panel of top SF3B1^MUT^ targets^45^. But since only a small number of transcripts were analyzed, we wished to examine a larger cohort of SF3B1^MUT^ targets. For this, we utilized our core list of 123 transcripts, and found that loss of DDX46 or DDX42 reproduced only 9.1% or 8.1% of SF3B1^MUT^-induced missplicing events, respectively (Figure 4*A, B*). Compared to AQR and SUGP1 loss, the effects of DDX46 and DDX42 loss on recapitulating SF3B1^MUT^-induced missplicing were very limited (Figure 4*D*), likely reflecting chance overlap and essentially identical to the overlap observed with all other SF/RBPs tested (Figure 4*D*). Additionally, we manually examined the raw RNA-seq data for several well-known SF3B1^MUT^ splicing targets, and neither DDX46 nor DDX42 loss caused any splicing defects (Figure 4*F*). These findings, together with those described above, highlight the unique role of SUGP1 in SF3B1^MUT^-induced missplicing.

**Fig. 4.**
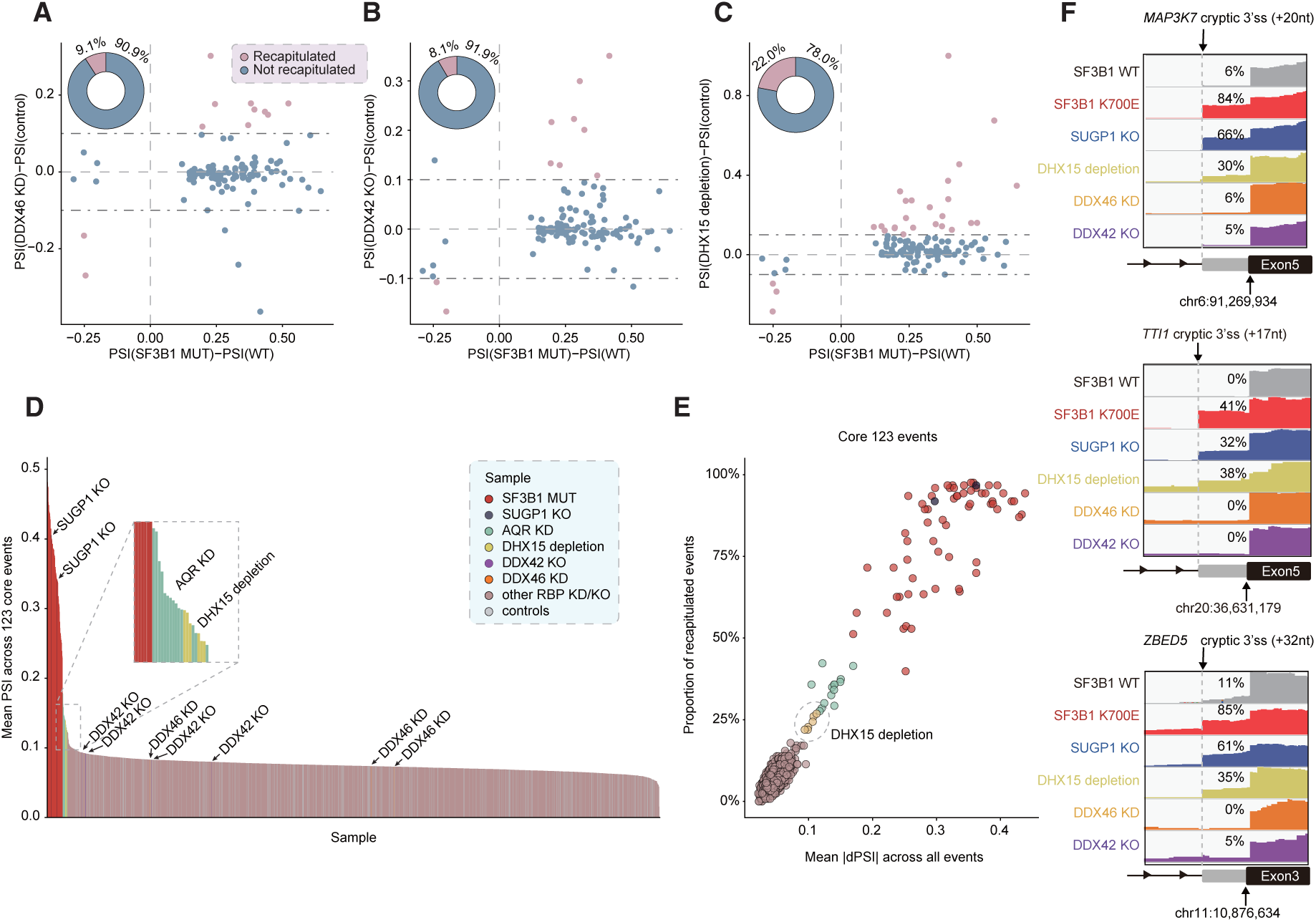
Loss of DHX15 but not DDX42/46 partially recapitulates SF3B1^MUT^-specific cryptic 3’ss usage. (*A*) Scatter plot representing the ΔPSI values for 123 core events in DDX46 KD samples compared with SF3B1^MUT^ samples. The horizontal axis shows the ΔPSI value between DDX46 KD and control samples, while the vertical axis shows the ΔPSI value between SF3B1^MUT^ sample and control sample. Pie plot (upper-left) indicates the proportion of 3’ss events recapitulated (brown) and not recapitulated (blue) by DDX46. (*B*) Similar to (*A*) but for DDX42. (*C*) Similar to (*A*) but for DHX15. (*D*) Each vertical bar represents the mean of the PSI values of the 123 core events in each of the SF/RBP loss and control samples. (*E*) Scatter plot representations of SF/RBPs whose expression loss positively correlated with cryptic 3′ss usage. The horizontal axis shows the average absolute ΔPSI value between RBP KD/KO samples and controls, and the vertical axis shows the percentage of recapitulated events. Analysis was performed on the 123 core events. (*F*) IGV plots of SF3B1^MUT^-specific cryptic 3’ss events for the indicated transcripts detected in WT, SF3B^K700E^, SUGP1 KO, DHX15 depletion, DDX46 KD and DDX42 KO. PSI values are shown for each track. DHX15 depletion was induced using degron tag technology^45^.

As mentioned above, previous studies have indicated that the DHX helicase DHX15 functions in concert with SUGP1 to ensure accurate BS selection in SF3B1^MUT^-target introns. However, DHX15 was not among the 600 SF/RBP KD/KO samples in the ENCODE and GEO databases we used for our analyses. We therefore examined whether DHX15 loss recapitulates SF3B1^MUT^-induced missplicing using six DHX15-depleted samples and controls from the study of Feng et al.^46^ (Table S10). Notably, DHX15 was very efficiently depleted in these experiments using conditional degron tags, rather than KD/KO. Using the same computational parameters as above, we found that DHX15 depletion recapitulated 22% to 27% of the missplicing events in our 123 event core list (Figure 4*C*, Tables S11, S12). Albeit not as robust as observed with SUGP1 or AQR loss (Figure 4*D-F*), these findings support the role of DHX15 in SF3B1^MUT^-induced missplicing, as suggested previously^23,24^.

## Discussion

In this study, we generated lists of high-confidence cryptic 3’ss events induced by *SF3B1* hotspot mutations. While other studies have identified SF3B1^MUT^ target transcripts^10–17^, our lists are unique in that they are comprehensive and rely on data from numerous studies involving a variety of cell and cancer types. These lists, which captured all previously studied cryptic 3’ss targeted by SF3B1^MUT^, including *MAP3K7*, *TMEM14C*, *ABCB7*, *ORAI2*, *PPOX* and *PPP2R5A*, thus represent a valuable resource to drive the biological and functional interrogation of cancer-specific mRNA isoforms. We used this new benchmark to perform an unbiased computational screen of 600 SF/RBPs to identify any whose loss, by KD or KO, recapitulates mutant SF3B1 splicing dysregulation. The results revealed that only loss of SUGP1, and to a lesser extent AQR, likely functioning indirectly, replicated the cryptic 3’ss use associated with *SF3B1* mutations, while a separate analysis identified DHX15, previously implicated in SUGP1 function. Below we discuss these findings and their significance with respect both to mechanisms necessary for correct BS recognition during splicing, as well as to how cancer-associated mutations affect this process and cause missplicing.

We have provided here strong evidence that loss of SUGP1 from pre-spliceosomal complexes is the primary, if not exclusive, cause of SF3B1^MUT^-induced missplicing in cells harboring *SF3B1* hotspot mutations. This conclusion is based on the nearly complete overlap between the cryptic 3’ ss activated by *SUGP1* KO and those used in SF3B1^MUT^ cells. Several previous studies are consistent with this result. Our initial study, which provided the first evidence that *SF3B1* cancer-associated hotspot mutations alter splicing by disrupting interaction with SUGP1, showed by RT-PCR that missplicing of a panel of SF3B1^MUT^ target transcripts could be recapitulated by SUGP1 KD^22^. Alsafadi et al., in their study identifying cancer-associated mutations in *SUGP1*, found by RNA-seq that SUGP1 KD recapitulated splicing defects brought about by overexpression of the SF3B1^K700E^ in HEK293 cells^20^. Similarly, Feng et al. ^46^, in a study providing evidence supporting a functional interaction between SUGP1 and DHX15, also observed a significant overlap in missplicing between SUGP1 KD cells and cells expressing SF3B1^K700E^. None of these studies, however, addressed the question of whether SUGP1 loss is exclusively responsible for SF3B1^MUT^ missplicing, as we have shown here.

As mentioned above, the RNA helicases DDX46 and DDX42 have also been suggested to function in SF3B1^MUT^-induced missplicing. The two helicases have been shown to compete for binding to the hotspot region of SF3B1 through their N-terminal domains during formation of the branch stem-loop of U2snRNA, allowing formation of the U2-BS RNA duplex^26–28^. Mutations altering the hotspot region in SF3B1, such as K700E, disrupt the interaction with DDX46/DDX42^28^. However, previous studies^16,20,21^ found no evidence that the genes encoding either of these two helicases harbor cancer-associated mutations that mimic the cryptic 3’ splicing caused by *SF3B1* mutation. This contrasts with the identification of such mutations in *SUGP1*^20,21^, the consequences of which are now understood mechanistically^24^. Moreover, as shown here, neither DDX46 KD nor DDX42 KO recapitulated *SF3B1* mutation-associated splicing dysregulation. Thus, not all proteins interacting with the SF3B1 hotspot region contribute to the aberrant BS selection resulting from *SF3B1* mutation. One explanation for this is that DDX46/DDX42 are recruited to interact with SF3B1 at a later stage in splicing, after discharge of SUGP1 from the spliceosome^24^. Once BS identification is complete, facilitated by SUGP1/DHX15, changes in DDX46/DDX42 levels will no longer impact BS recognition^47^.

Previous studies provided conflicting results concerning the role of DHX15 in mediating the effects of mutant SF3B1. Our study that established the interaction between SUGP1 and DHX15 revealed an ∼13% overlap between SF3B1^K700E^ and DHX15 KD samples^23^, while Feng et al. reported no significant overlap between their DHX15 degron-depleted samples and another unspecified SF3B1^K700E^ sample^46^. Why these studies gave disparate results, and differed from those reported here, is unclear, but likely reflects the nature and size of the datasets analyzed. In any case, while the SF3B1^MUT^-like missplicing we detected here in the DHX15-depleted samples was greater than observed with any other SF/RBP except SUGP1 or AQR, an interesting question is why DHX15 loss did not result in the extensive missplicing caused by SUGP1 depletion. While additional studies will be required to address this, it could reflect the fact that DHX15 is involved in several cellular processes^44^ and can interact with multiple G-patch proteins^48^, thereby, as we discussed previously^23^, possibly suppressing certain SUGP1-dependent missplicing events. Another possibility is that a different DHX helicase might partially compensate for DHX15 loss.

It was initially surprising that AQR was the only protein other than SUGP1, and to a lesser degree DHX15, whose loss significantly recapitulated SF3B1^MUT^-induced missplicing. AQR has been shown to be incorporated into spliceosomes as part of a pentameric intron-binding complex that associates with U2snRNP and is essential for splicing in vitro^33^. However, based on cryo-EM and biochemical data, AQR appears to function after BS recognition, facilitating transfer of the BS-U2snRNA hybrid to the catalytic center of the spliceosome and release of SF3A and B complexes from the spliceosome^35^. Thus, a direct role for AQR in activation of cryptic 3’ss in SF3B1^MUT^ cells seemed unlikely, and our finding that *SUGP1* pre-mRNA is among the many thousands of misspliced transcripts in AQR-depleted cells provides a parsimonious explanation for the SF3B1^MUT^-like missplicing found in these cells. Indeed, the 50–60% reduction of SUGP1 protein upon AQR KD is consistent with the fact that only ∼40% of cryptic 3’ss events induced by SF3B1^MUT^ were recapitulated by AQR KD. It is also notable that although AQR loss leads to SUGP1 downregulation by inducing use of a cryptic 3’ss in *SUGP1* pre-mRNA, this site is not used in SF3B1^MUT^ cells. This finding indicates that AQR loss induces missplicing by a different mechanism than do *SF3B1* cancer-associated mutations, consistent with the structural studies noted above^49^.

In summary, our study has provided a comprehensive list of core splicing events that are dysregulated by cancer-driving mutations in *SF3B1* in multiple different cancer types, which should be of considerable value in elucidating the consequences of such mutations. Using this list, we found that among hundreds of splicing-related proteins, KD or KO of only two, SUGP1 and AQR, replicated the missplicing events induced by mutant SF3B1. However, because AQR likely functions indirectly, by reducing SUGP1 expression, our findings indicate that SUGP1 loss from mutant spliceosomes is the sole driver of cancer-associated splicing dysregulation induced by *SF3B1* mutations.

## Methods

### Pan-cancer SF3B1-mutated sample collection

We collected RNA sequencing data from both patient and cell line samples with and without SF3B1 hotspot mutations from the TCGA (https://portal.gdc.cancer.gov) and GEO databases (https://www.ncbi.nlm.nih.gov/geo). In total, we collected 63 SF3B1 mutant samples and 56 WT samples, encompassing four tumor types (MDS, CLL, UVM and SKCM) and two cell lines (K562 and Nalm6). Detailed sample information is provided in Table S1.

### RBP KO/KD sample collection

All RNA-seq data for RBP KD and KO samples (including shRNA, siRNA, CRISPR and CRISPRi RNA-seq) were downloaded from the ENCODE project phase III database (https://www.encodeproject.org/publications/4928df3e-6995-4401-b643-84980bb94057/) (24). Additionally, we collected RNA-seq data for 187 KD/KO samples targeting 35 RBPs from the GEO database. These RBPs were identified as physically interacting with SF3B1 using mass spectrometry in our previous work^22^. In total, we compiled 2,151 RBP-related KD or KO samples and 298 matched control samples, covering 600 RBPs and 18 cell types. Detailed sample information is provided in Table S4.

### Identification of cryptic 3’ss events

We designed a computational analysis pipeline to identify and quantify the usage of all annotated and novel cryptic splice junctions, as well as canonical 3’ss associated with cryptic 3’ ss. Briefly, FASTQ files of the RNA-seq data for SF3B1^MUT^ and SF3B1^WT^ sample were aligned to human genome (hg19) using STAR version 2.7.11a, with the reference genome annotation GTF file downloaded from GENCODE database in release 19 (https://www.gencodegenes.org/human/release_19.html). Then, junction reads counts from the STAR output file (SJ.out.tab) was merged into a matrix, with each row representing a splice junction and each column indicating a sample. Low-abundance junctions with fewer than 200 reads (summed up across all samples) were filtered out. For the remaining junctions, we compared each one to known splice junctions in the reference GTF file. A junction was defined as alternatively spliced if there is another junction shared the same annotated 3’ or 5’ end. The associated canonical junction was then identified if it shared the same annotated 3’ or 5’ end with the cryptic splice junction and had the maximum supporting reads in SF3B1^WT^ samples. We then determined the relative position of each alternative 3’ or 5’ splice site to the associated canonical 3’ or 5’ splice site based on their locations on the same transcript strand. Cryptic splicing events where the distance was less than or equal to 3 base pairs (bp) or greater than 100 bp were filtered out. Next, we calculated the PSI values from the raw junction read counts. We performed t-tests to obtain *p*-values by comparing the PSI values between SF3B1^MUT^ and SF3B1^WT^ samples across six SF3B1^MUT^-cohorts. Additionally, we calculated ΔPSI, defined as the difference between the mean PSI of SF3B1^MUT^ and SF3B1^WT^, across six cohorts. Finally, after manual verification by IGV, we retained only those cryptic 3’ss events with a *p*-value <0.05 and an absolute ΔPSI >0.1 in more than 2 out of 6 cohorts, and defined this set as the SF3B1^MUT^-specific events (295 list). Additionally, cryptic 3’ss events with a *p*-value <0.05 in more than 2 cohorts and an absolute ΔPSI >0.1 in all 6 cohorts were defined as the core list of events induced by SF3B1 mutant (123 list). (See also in Figure 1A.)

For cryptic 3’ss events induced by SUGP1 loss, we applied the same workflow on SUGP1 KO RNA-seq data from GSE242092 and control RNA-seq data from GSE187356, with a total junction read count threshold of 10. After calculating ΔPSI and *p*-value between the SUGP1 KO and control samples, we sorted the cryptic 3’ss events by *p*-value and retained the top 295 cryptic 3’ss events specific to SUGP1 loss.

### Evaluation of effects of RBP loss on recapitulating SF3B1^MUT^ specific cryptic 3’ss events

RBP KD and KO samples were processed by aligning the FASTQ files to human genome (hg19) using STAR version 2.7.11a, employing the reference genome annotation GTF file downloaded from GENCODE (https://www.gencodegenes.org/human/release_19.html). Next, for each sample, we extracted alternative junction read counts and canonical junction read counts of 295 SF3B1^MUT^-specific cryptic 3’ss events from corresponding SJ.out.tab to calculate the PSI value (PSI^RBP^). For each of these 295 cryptic 3’ss events, we summed the alterative and canonical junction reads, respectively, across all control samples to establish the background PSI value (PSI^Con^). Then, a cryptic 3’ss event was deemed to be recapitulated by RBP KD/KO if the absolute value of dPSI (PSI^RBP^-PSI^Con^) ≥0.1 and the event exhibits the same splicing direction in RBP KD/KO and SF3B1^MUT^. Lastly, all RBP KD/KO samples were visualized based on the proportion of recapitulated events and mean absolute dPSI (|PSI^RBP^-PSI^Con^|).

### Identification the sequence characteristics of SF3B1^MUT^-specific cryptic 3’ss events

To identify the sequence characteristics of SF3B1^MUT^**-**specific cryptic 3’ss events, we used bedtools2.28 to obtain the nucleotide sequences 50bp upstream and downstream of both the cryptic and associated canonical 3’splice site from our list of 295 events. As a control, we selected 500 introns where no cryptic 3ss usage was detected (*p*-value = 1 between SF3B1 mutant and WT). For both the aberrant splicing sequences and the control sequences, we calculated the nucleotide composition of adenine (A), thymine (T), guanine (G), and cytosine (C) at each position. We then employed Fisher’s exact test to determine the significance of enrichment for each nucleotide in the aberrant group compared to the control group. Finally, we visualized the nucleotide percentages at each position using the R package ‘ggseqlogo’.

### Heatmap and hierarchical clustering

To explore the potential similarity of SF3B1^MUT^-specific cryptic 3’ss events alteration between SF3B1 mutants and different RBP KD/KO, principal component analysis and unsupervised hierarchical clustering were performed using 295/123 alternative 3’ss event induced by SF3B1^MUT^. We utilized the ‘prcomp’ command from the R package ‘stats’ for principal component analysis. For clustering analysis, we employed the ‘Heatmap’ command from the R package ‘ComplexHeatmap’, using Euclidean distance as the metric.

### Identification of alternative splicing events induced by SRSF2^MUT^ and U2AF1^MUT^

For SRSF2^50^ FASTQ files were downloaded from the GEO database via access id GSE71299, and aligned to the human genome (hg19) using STAR version 2.7.11a, with reference genome annotation GTF file downloaded from GENCODE database in release 19 (https://www.gencodegenes.org/human/release_19.html). Then, rMATs (version 4.1.1) was used to identify the differential exon skipping (ES) events between SRSF2 mutant and WT (4 vs 4). We extracted the inclusion level (IncLevel) values from the [Junction Counts and Exon Counts] (JCEC) file to create a matrix with IncLevel as the variable across samples. Subsequently, we recalculated the *p*-values using a t-test between SRSF2^MUT^ and WT to determine the inclusion level difference. Events were filtered by a *p*-value of 0.05 and an inclusion level difference of 10%. Additionally, we collected SRSF2^MUT^-specific events that were experimentally validated from the ASCancer Atlas database (https://ngdc.cncb.ac.cn/ascancer/home). After manual confirmation using IGV, 42 events were retained. Finally, by sorting based on *p*-values and including the 42 experimentally validated events, a total of 295 SRSF2^MUT^-specific AS events were obtained (See also Figure S2*D*). For U2AF1, we applied the same workflow as above on three cases of U2AF1^MUT^ and three cases of U2AF1^WT^ RNA-seq data to obtain 295 U2AF1^MUT^ specific AS events. (See also Figure S2*E*.)

### Cell lines

CRISPR-engineered K562 cells with WT or K700E mutant SF3B1 (three clones each) were generated in our previous study^22^. These CRISPR cells and parental K562 were cultured in Iscove’s Modified Dulbecco’s Medium (IMDM; Gibco, cat # 12440-053) supplemented with 10% fetal bovine serum (FBS; Seradigm, cat # 89510-186) in a 37 °C, 5% CO2 incubator.

### Knockdown experiments

K562 cells were electroporated with two independent AQR siRNAs using the Neon Transfection System (Thermo Fisher Sci) with the Neon 100uL tip kit that includes the R buffer for electroporation (Thermo Fisher Sci). One million cells were resuspended in 100uL of R buffer, containing 1uL of 100uM siRNAs. Transfection was performed following the manufacturer’s protocol. Electroporated cells were immediately placed in 2mL complete media for 24h for a final siRNA concentration of 50uM. After 24h, cells were washed with PBS and electroporated again with the same siRNAs (second round) and placed in 2ml of complete media for another 24h at a 50uM concentration. Cells were harvested at 48h after the second round of electroporation for downstream assays. The sequences of the two siRNAs targeting AQR were: siAQR-1 sense strand (5′-GAGAUUGUCAAAUCAAGGUdTdT- 3′), siAQR-1 antisense strand (5′-ACCUUGAUUUGACAAUCUCdTdT-3′), siAQR-2 sense strand (5′-GGCGCUGGUUUAAUACCAUdTdT-3′), and siAQR-2 antisense strand (5′-AUGGUAUUAAACCAGCGCCdTdT-3′). The sequences of the sense and antisense strands of the negative control siRNA (siC) were listed in our previous study^22^. For emetine treatment, a final concentration of 100 μg/ml emetine dihydrochloride was added to the cells at 36h post-second-round siRNA electroporation. After 12h of incubation, the cells were harvested for RNA extraction.

### Western blotting

Western blotting was performed as described^22^ with the following primary antibodies: anti-AQR (1:1,000, Bethyl Laboratories, A302-547A-T), anti-SUGP1 (1:1,000, Sigma-Aldrich, HPA004890), anti-ACTIN (1:2,000, Sigma-Aldrich, A2066), anti-SETX (1:1000, Bethyl Laboratories, A301-105A). Secondary antibodies used were Goat anti-Rabbit IgG (H+L) Secondary Antibody, HRP (ThermoFisher, cat #31460) and Goat anti-Mouse IgM (Heavy chain) Secondary Antibody, HRP (ThermoFisher, cat# 62-6820). Membranes were detected using the Pierce™ ECL Western Blotting Substrate chemiluminescent reagent (ThermoFisher cat# 1859698). The fluorescence secondary antibody used was Donkey anti-Rabbit IgG (LI-COR, 926-68073, 1:5,000). Fluorescence signals were detected using the ChemiDoc Imaging System (Bio-Rad).

### Statistical analysis of western blots

Chemiluminescent/fluorescence signals were analyzed using ImageJ and the band intensity of the target proteins (SUGP1, AQR and SETX) was divided by the band intensity of the housekeeping proteins used (ACTB and TUB). The ratios obtained from siAQR-1 and siAQR-2 were then normalized to the ratios of siControl samples. The normalized ratios from 3 independent experiments were then analyzed using an ordinary one-way ANOVA with Dunnet’s multiple comparison test wherein the mean of each columns (siAQR-1 and siAQR-2) was compared to the mean of a control column (siC).

### Reverse Transcription-Polymerase Chain Reaction

Total RNA was extracted with TRIzol (Thermo Fisher Scientific). In a 20-ul reaction, 2 μg total RNA was reverse-transcribed using 0.3 μl Maxima Reverse Transcriptase (Thermo Fisher Scientific) and 50 pmol oligo-dT primer. The cDNA was diluted (1:10) and 3 μl used for PCR. PCR products were subjected to agarose gel electrophoresis with ethidium bromide staining, followed by gel imaging with the ChemiDoc Imaging System (Bio-Rad). Primers used were *MED6* forward, 5′- CTGGAGTGATCTATCAGGCACC-3′; *MED6*reverse, 5′- GCCACCAATACCCTTTGGAAGG-3′; *TOR1AIP2* forward, 5′- GTTGGTCTCGAACTCCTGGCTT-3′; *TOR1AIP2* reverse 5′- TGAAGAGTCTGGGAATGCGTCAC-3′; *WASHC5* forward, 5′- GGGCAGATGCAGATTCTGAGAC-3′; *WASHC5* reverse, 5′- GTGAAGGGTCCTGATAGTGGG-3′; *ZNF410* forward, 5′- ACAGGAAGGTATCATTGGCTCTG-3′; *ZNF410* reverse, 5′- TCTACAAGCCCCTGTCCCAATG-3′; *SUGP1* forward, 5′- AAAGGAAGCACAGAAGTCGCAG-3′; and *SUGP1* reverse, 5′- TTCTCACGGTTGTTCTGGAGGG-3′. Primers for *GCC2* and *MAP3K7* were listed in our previous study^22^.

## Data availability

All possessed data generated in this study are included in the main text, SI Appendix, or Datasets S1–S11. The RNA-sequencing data of U2AF1 mutant and matched WT samples have been deposited in NCBI’s Gene Expression Omnibus^51^ and are accessible through GEO Series accession number GSE285785 (https://www.ncbi.nlm.nih.gov/geo/query/acc.cgi?acc= GSE285785).

## Acknowledgments

This work was supported by the Beijing Natural Science Foundation (grant Z220012 to Z.L.), the National Natural Science Foundation of China (grant 32170565 to Z.L.), the Chinese Academy of Sciences Hundred Talents Program (to Z.L.). This work was also supported by NIH grant R35 GM118136 to J.L.M. We thank Dr. Yen Lieu for her help during the early stages of this work.

## Supplemental Figure Legends

**Fig. S1.**
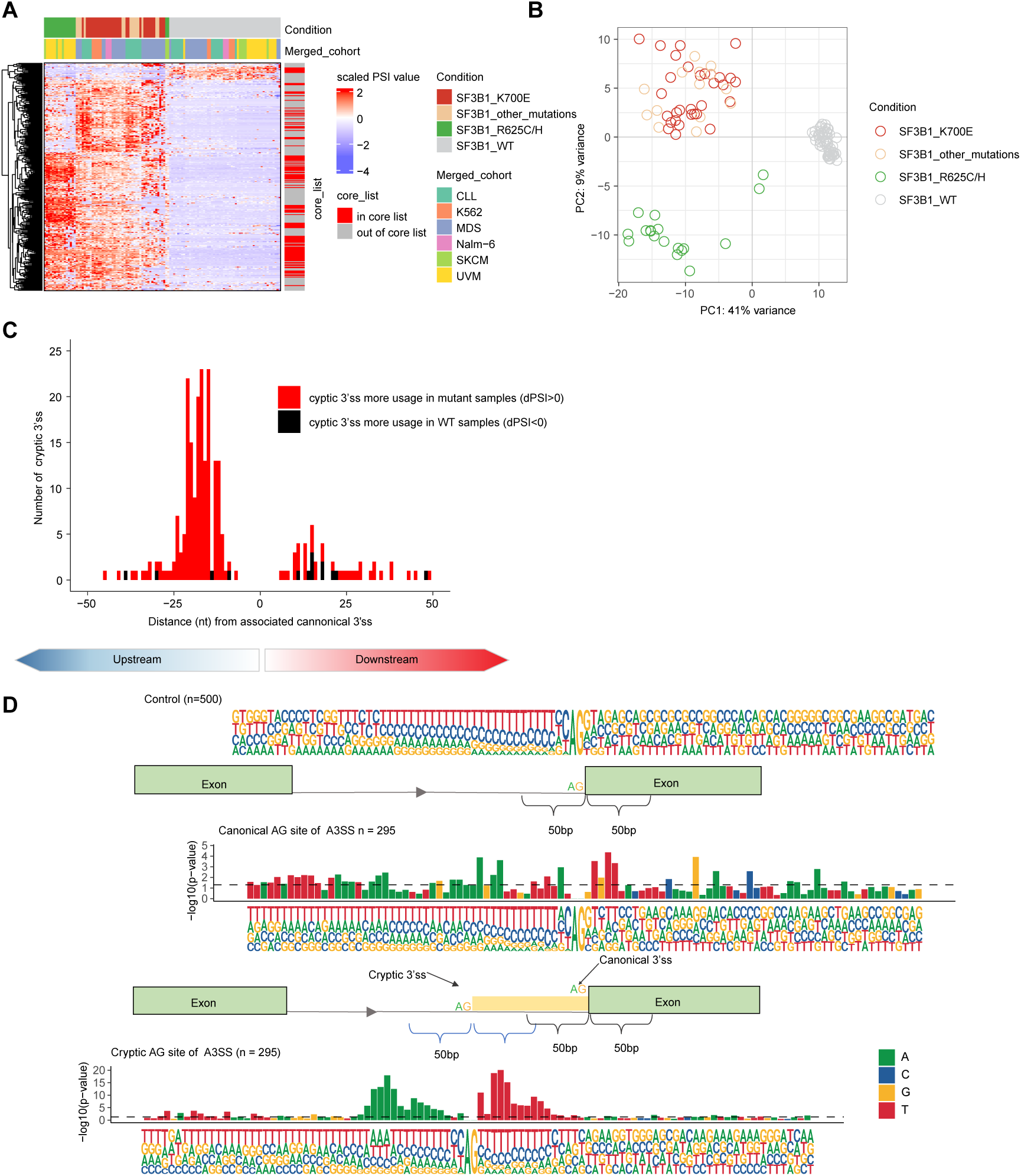
Characteristics of SF3B1^MUT^-specific cryptic 3’ splicing events. (*A*) Hierarchical clustering using Euclidean distance coupled with heatmap analysis illustrates usage of 295 3′ss in all SF3B1^MUT^ and WT samples. Rows represent the 295 cryptic 3′ss events, while columns represent the samples. Values in the matrix correspond to row-scaled normalized PSI. (*B*) Principal component analysis (PCA) of PSI values of all SF3B1^MUT^ and WT samples focusing on the 295 cryptic 3’ss events. (*C*) Density plot of distance (in bp) from associated canonical 3’ss to cryptic 3’ss. (*D*) Consensus 3’ss motif near the associated canonical 3’ AG dinucleotide (middle track) and cryptic 3’ AG dinucleotide (bottom track). Top track displays sequence motif of 500 control events in which cryptic 3’ss usage was not detected. The sequence range shown is 50nt upstream and 50nt downstream of the AG site. The size of a single “A G T C” represents the frequency of occurrence at each position. Bar plots above each nucleotide composition plot are –log10(*p*-value) from Fisher exact tests for enrichment of nucleotides at each position relative to controls. Horizontal dash line marks significance level of *p*-values =0.05 (see details in Methods).

**Fig. S2.**
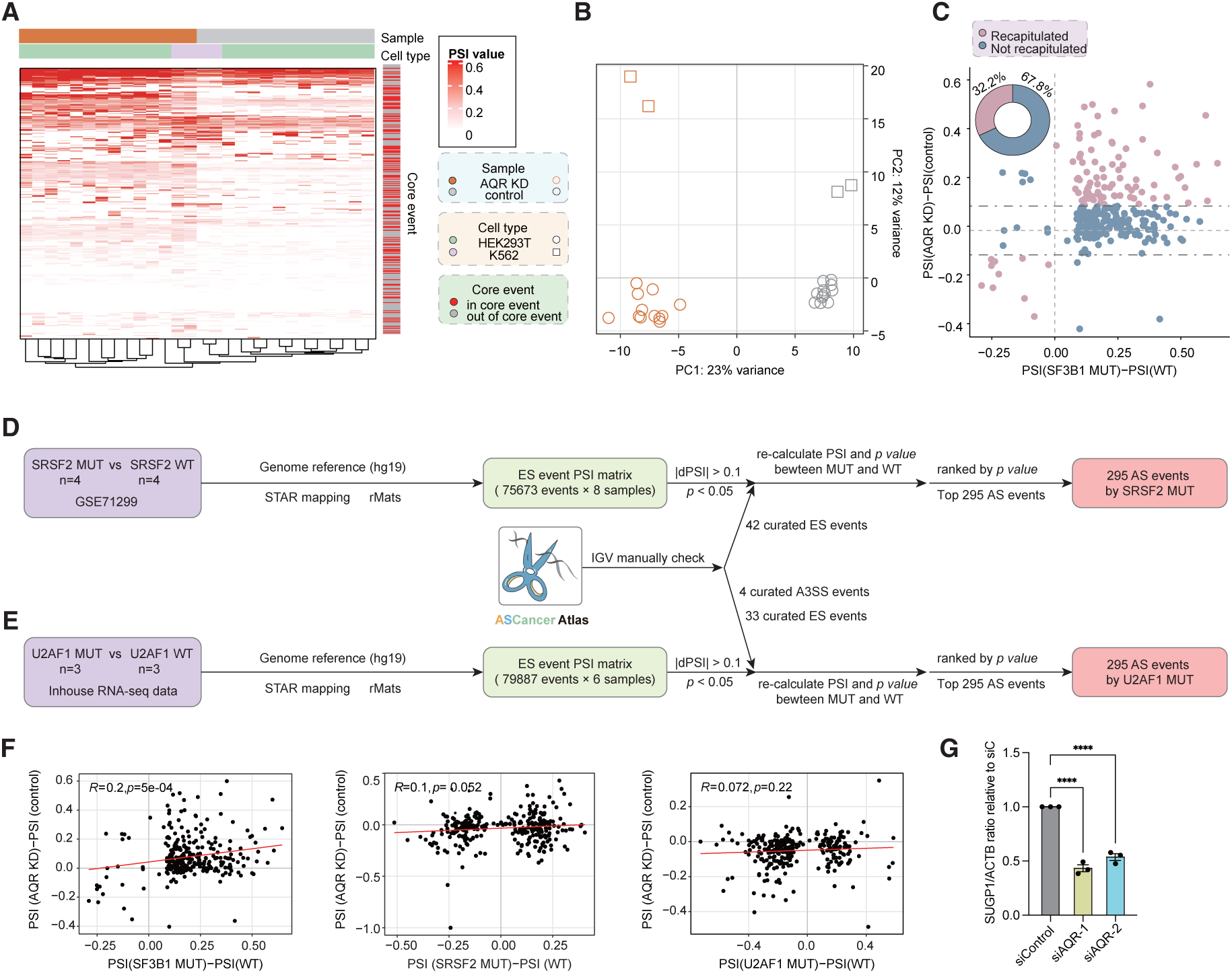
AQR loss recapitulates ∼40% SF3B1^MUT^-specific cryptic 3’ss events reflecting SUGP1 missplicing. (*A*) Hierarchical clustering and heatmap analysis of the usage of 295 SF3B1^MUT^-specific events in AQR KD samples and corresponding control samples. Rows represent the 295 SF3B1^MUT^-specific events, while columns represent the samples. The two-column annotation bar plot at the top of the heatmap indicates the experimental group and cell type. (*B*) PCA of PSI value of all AQR KD and corresponding control samples with 295 SF3B1^MUT^-specific events. (*C*) Scatter plot representing the ΔPSI values for 295 SF3B1^MUT^-specific events in AQR KD compared with SF3B1^MUT^ samples. The horizontal axis shows the ΔPSI value between SF3B1^MUT^ and control samples, while the vertical axis shows the ΔPSI value between AQR KD sample and control sample. Pie plot (upper-left) indicates proportion of cryptic 3’ss events recapitulated (brown) and not recapitulated (blue) by AQR KD. (*D*) Workflow to identify SRSF2^MUT^-specific AS events. (*E*) Similar to (*D*) but for U2AF1. (*F*) Scatter plot indicating the impact of AQR KD on specific events in SF3B1^MUT^, SRSF2^MUT^ and U2AF1^MUT^ samples. The horizontal axes shows the PSI changes comparing SF3B1^MUT^, SRSF2^MUT^ and U2AF1^MUT^ samples to WT samples, while the vertical axis shows the PSI changes comparing AQR KD to control samples. R values and *p*-values were determined by Pearson correlation. (*G*) Quantification of the SUGP1/ACTB ratio relative to siC in Figure 2D. Bars represent the mean ± SEM (n = 3; three independent experiments). Ordinary one-way ANOVA with Dunnet’s multiple comparison test wherein the mean of each columns (siAQR-1 and siAQR-2) was compared to the mean of a control column (siC). \**p* <0.05, \*\**p* <0.01, \*\*\**p* <0.001 and \*\*\*\**p* <0.0001.

**Fig. S3.**
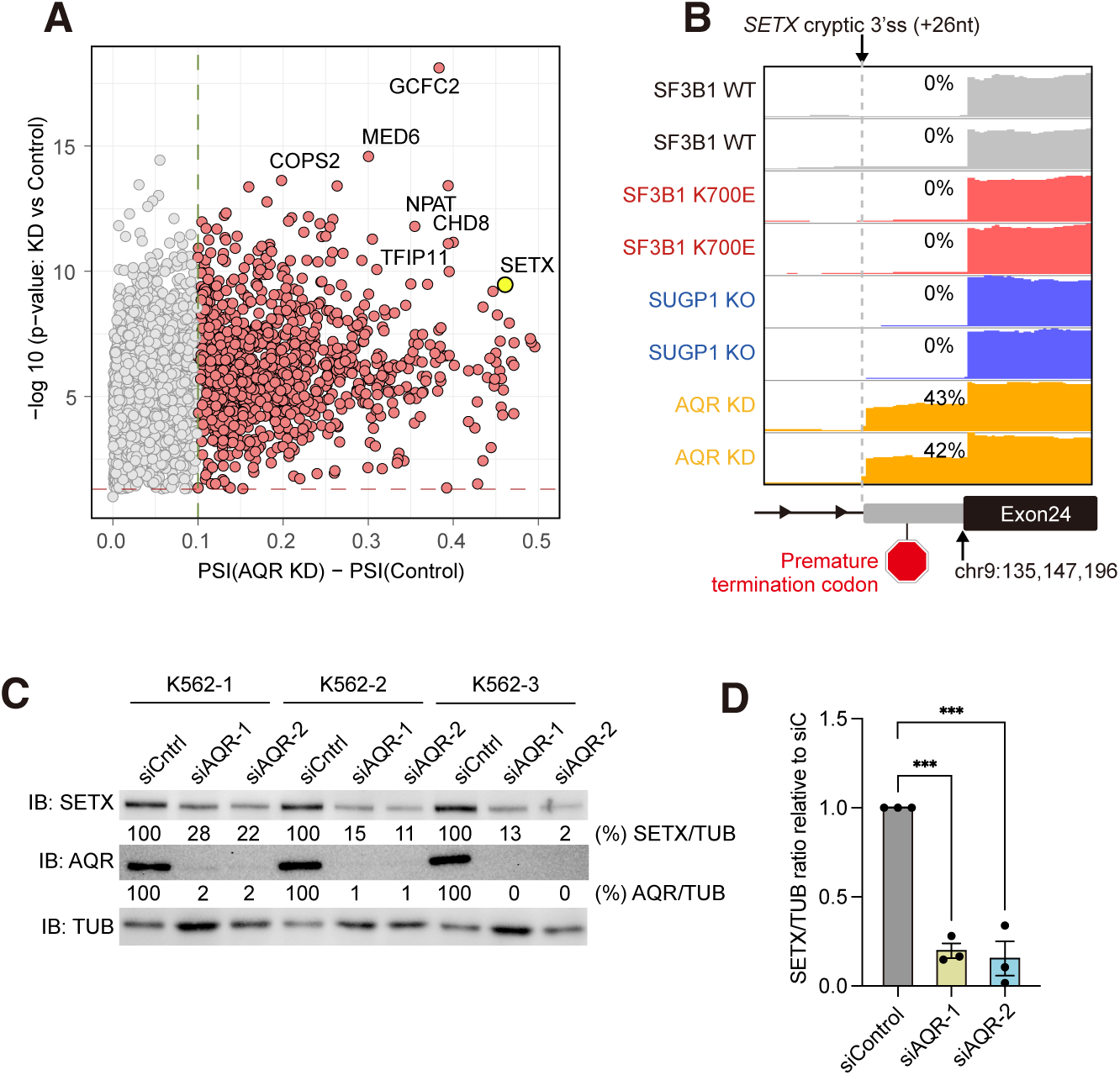
AQR KD causes missplicing of *SETX* transcripts and reduced protein levels. (*A*) Scatter plot representation of differentially spliced 3’ss on all transcripts between AQR KD and controls showing the magnitude (difference of PSI; x-axis) and significance (-log10(q-value); y-axis). Identities of select transcripts are indicated. (*B*) IGV plots of the cryptic 3’ss event in SETX intron 23 in WT, SF3B1^K700E^, SUGP1 KO and AQR KD samples. PSI values are shown for each track. (*C*) Western blot analysis of SETX, AQR and TUB in extracts from AQR KD K562 cells. SETX/TUB and AQR/TUB ratios from three independent experiments are highlighted, which were used for the statistical analysis (see Methods). (*D*) Quantification of the SETX/TUB ratio relative to siC from WB in (*C*). Bars represent the mean ± SEM (n=3; three independent experiments). Ordinary one-way ANOVA with Dunnet’s multiple comparison test wherein the mean of each column (siAQR-1 and siAQR-2) was compared to the mean of a control column (siC). \**p* <0.05, \*\**p* <0.01, \*\*\**p* <0.001 and \*\*\*\**p* <0.0001.

**Fig. S4.**
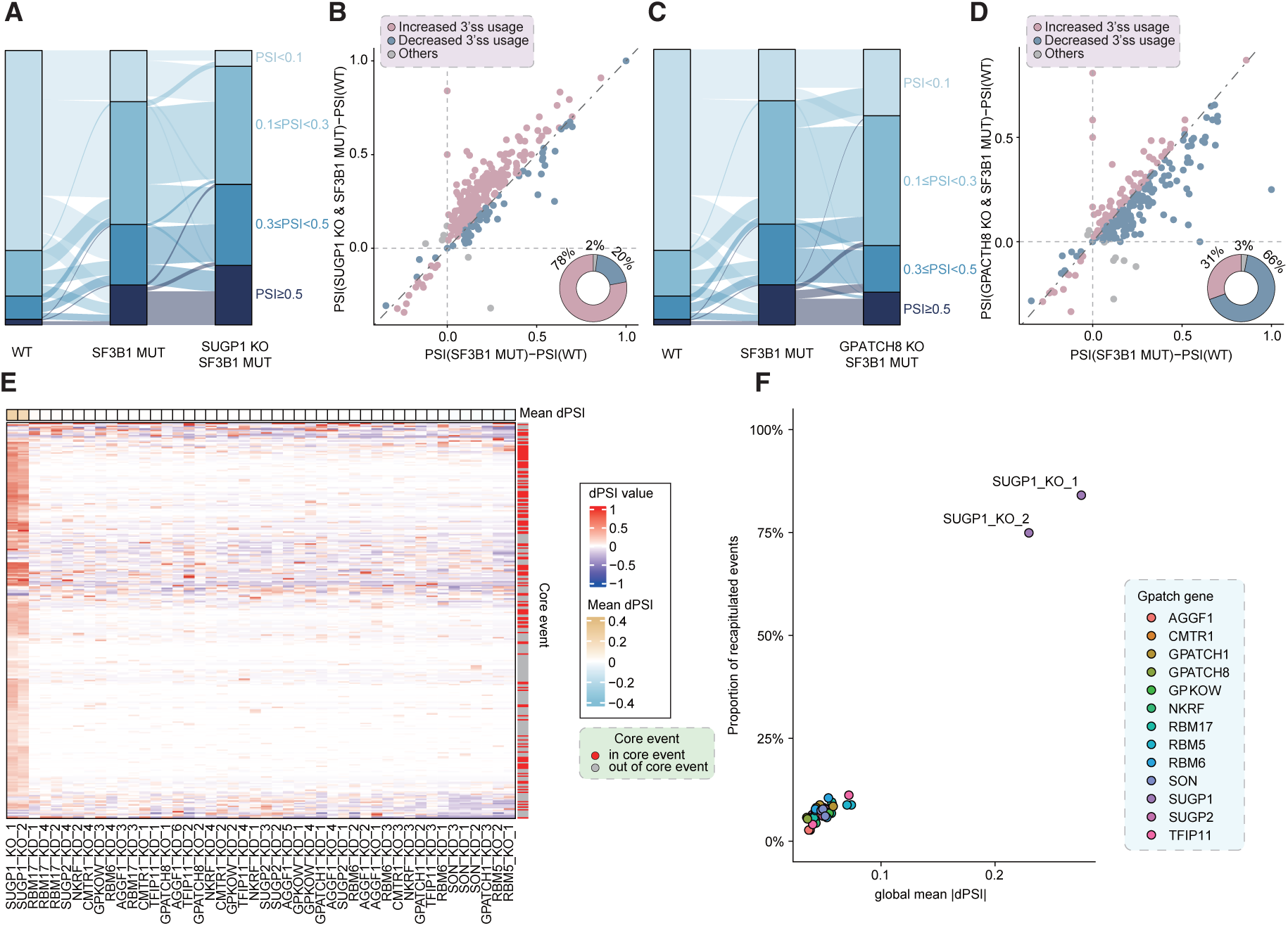
Distinct effects of G-patch protein loss on SF3B1^MUT^-specific cryptic 3’ss events. (*A*) Sankey diagram illustrating changes in PSI values of 295 SF3B1^MUT^-specific events due to *SF3B1* mutation and the combined effect of *SF3B1* mutation with SUGP1 KO. Different colors represent the range of PSI values across different groups. (*B*) Scatter plot depicts ΔPSI values for 295 SF3B1^MUT^-specific events in SF3B1^MUT^ cells compared with SF3B1^MUT^ and SUGP1 KO cells. The horizontal axis shows the ΔPSI values between SF3B1^MUT^ and control samples, while the vertical axis shows the ΔPSI value between SF3B1^MUT^ plus SUGP1 KO sample and control sample. Pie plot (lower-right) indicates the proportion of increasingly (brown) and decreasingly (blue) used cryptic 3’ss in the SF3B1^MUT^ and SUGP1 KO samples compared with the SF3B1^MUT^ samples. (*C*) Similar to (*A*) but for GPATCH8. (*D*) Similar to (*B*) but for GPATCH8. (*E*) Hierarchical clustering and heatmap analysis of the ΔPSI value of 295 SF3B1^MUT^-specific events in G patch domain-containing protein (G-patch gene) loss samples. Rows represent the 295 SF3B1^MUT^-specific events, while columns represent the samples. The column annotation bar plot at the top of the heatmap indicates the average ΔPSI value across the 295 SF3B1^MUT^-specific events. (*F*) Scatter plot representations of effects of G-patch protein loss on 295 SF3B1^MUT^-specific cryptic 3’ss events. The horizontal axis shows the average ΔPSI values between G-patch protein KD/KO and control samples, and the vertical axis shows the percentage of recapitulated events.

